# Structural basis of Cas12a R-loop propagation on pathway to DNA cleavage

**DOI:** 10.1101/2023.03.13.532460

**Authors:** Isabel Strohkendl, Catherine Moy, Alexander-Hoi Nguyen, Rick Russell, David W. Taylor

## Abstract

Cas12a is a CRISPR RNA-guided nuclease that engages target DNA through protein-DNA and RNA-DNA contacts. Initial PAM binding by Cas12a leads to formation of a 20bp R-loop between the complementary crRNA guide and target strand. Following specificity-determining R-loop formation, both DNA strands undergo RuvC-mediated cleavage. Current structures of Cas12a bound to its target only show the R-loop after formation, leaving an important gap in knowledge as to how Cas12a accommodates the extending R-loop and coordinates R-loop formation with nuclease activation. Here, we use cryo-EM to capture a series of kinetically trapped Cas12a R-loop intermediates and observe how Cas12a delivers each DNA strand into the RuvC active site. We show that Cas12a first interrogates target DNA via a 5bp seed, followed by dramatic Rec domain conformational flexibility to accommodate R-loop extension. Only during formation of the final R-loop base pairs do the Rec and BH domains engage in the majority of contacts with the R-loop. R-loop completion leads the nontarget strand to displace the RuvC lid and kink into the active site via a base stacking interaction. Following nontarget strand cleavage, we observe substantial Rec2 and Nuc domain dynamics as the TS is brought to the RuvC active site. Our kinetics-guided structural snapshots provide a comprehensive model describing Cas12a DNA targeting and highlight mechanistic differences between Cas12a and Cas9.

## Introduction

Cas12a has been repurposed for genome editing and biotechnological applications due to its readily adaptable RNA-guided targeting capabilities^1^. Cas12a specifically binds to unique DNA sequences, cleaves double-stranded DNA targets in *cis*, and nonspecifically cleaves ssDNA in *trans*^*2-5*^. After Cas12a assembles with and processes its crRNA, it scans DNA in search of a short, T-rich protospacer adjacent motif (PAM) to initiate target recognition. PAM recognition leads to local DNA melting that allows the crRNA guide to invade the duplex DNA and form a 20-base pair (bp) R-loop with the complementary target strand of the matched ‘on-target’ DNA^6^. R-loop formation triggers activation of the sole RuvC nuclease domain, creating a double stranded DNA break. On-target R-loop formation and cleavage occur in rapid succession, so mismatches that occur between the crRNA guide and target strand are kinetically discriminated against during R-loop formation and ensure high specificity in binding and cleavage^7^.

Cas12a functions similarly to Cas9 as both rely on rate-limiting R-loop formation to activate target DNA cleavage^8-10^. However, differences in activity and specificity profiles suggest they have distinct underlying mechanisms influenced by their protein architecture^11-13^. Cas12a undergoes large conformational changes during assembly into its binary and ternary complexes^14-19^. While it is known that the Rec lobe must rearrange to accommodate the helical 20bp R-loop and expose the RuvC nuclease domain, there is a lack of kinetically relevant structures along the reaction pathway that inform how these conformational changes are coordinated with R-loop propagation. Given the important role R-loop formation has in nuclease activation and off-target rejection, structural insights into Cas12a R-loop formation are required to understand its activity and specificity to guide future strategies for re-engineering the enzyme.

To understand how Cas12a interrogates its DNA target and transitions into the catalytically active state, we used cryo-electron microscopy (cryo-EM) to capture structures of wild type Cas12a R-loop formation on pathway to DNA cleavage. We show that Cas12a first interrogates 5 base pairs of target DNA before dramatic Rec domain reorganization accommodates R-loop extension. Only during formation of the final few base pairs does the Rec lobe fully dock onto the R-loop, forming protein contacts along the majority of the RNA-DNA heteroduplex via Rec1, Rec2, and bridge helix (BH) domains. Formation of the final base pairs also leads to the nontarget strand (NTS) traversing the RuvC domain and displacing the inhibitory RuvC lid. Kinetics-guided cryo-EM enabled capturing the NTS poised for cleavage in the RuvC active site, uncovering the mechanism of RuvC-mediated cleavage. We then captured Cas12a after NTS cleavage to observe dramatic Cas12a dynamics as the Rec and Nuc domains manipulate the downstream target strand (TS) to reach the RuvC active site. These structures help provide a detailed model that describes Cas12a DNA targeting and underscores mechanistic differences between Cas12a and Cas9.

### Kinetics-guided cryo-EM maps of Cas12a on pathway to cleavage

To capture various intermediates of Cas12a during R-loop formation on pathway to DNA cleavage, we designed DNA duplex substrates with 8, 12, 16, and 20 bp of complementarity to the crRNA guide assembled with WT *Acidaminococcus sp*. Cas12a (AsCas12a) (Table S2). Cas12a-crRNA-DNA binding incubation times varied to maximize binding with minimal cleavage (8 bp, 1 hr; 12 bp, 1 hr; 16 bp, 4 min; 20 bp, 1 min at ∼18° C (ambient temperature) before vitrification. From these four samples, we obtained six distinct R-loop intermediate structures with nominal resolutions ranging from 3.3-3.6 Å (Fig. 1, Fig. S1-5, Table S1). From these structures, we observed Cas12a R-loop formation in real time and noted several intriguing results: (1) the number of R-loop base pairs resolved is not always equal to the extent of target complementarity; (2) Rec2 is not resolved during R-loop propagation; and (3) the distal DNA migrates from projecting out the front of Cas12a to the back.

**Fig. 1.**
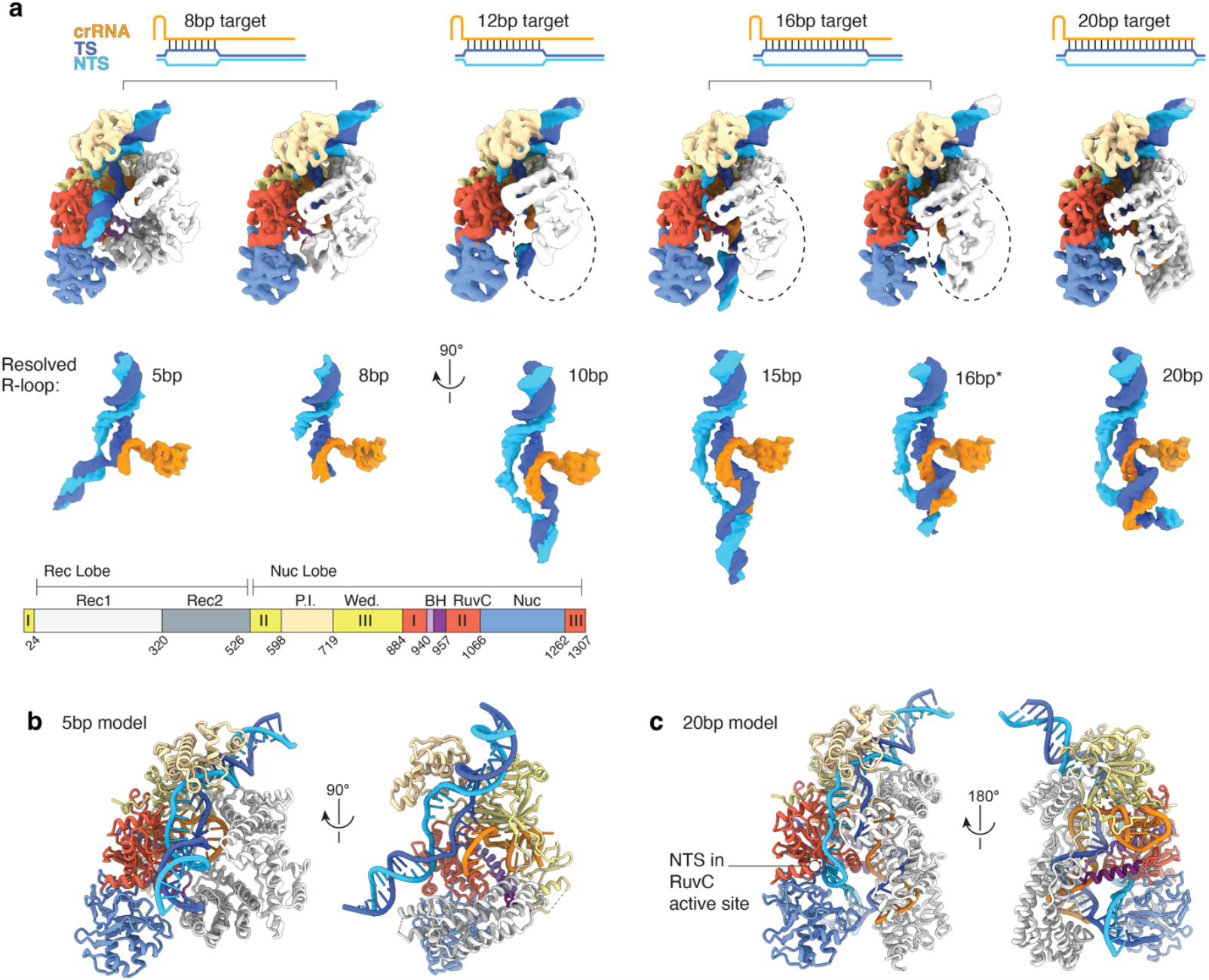
Cas12a R-loop intermediates captured by kinetics-guided cryo-EM. **a**, DNA-bound Cas12a cryo-EM structures with varying lengths of the R-loop resolved. Cryo-EM maps are grouped by DNA target used and ordered by resolved R-loop base pairs. Densities for nucleic acids are shown at a 90° angle to the overall structure. Varying thresholds were used to increase length of visible NTS. Maps are colored according to the domain map. **b**, Model of 5bp reconstruction. The side view has Rec1 removed to better view the seed R-loop. **c**, Model of the 20bp reconstruction with the NTS poised for cleavage..

The 8bp target dataset yielded two similarly populated structures with well resolved 5bp and 8bp of contiguous R-loop density (Fig. S2), revealing early contacts in target search and R-loop seeding. Further classification demonstrates the extent of Rec1 domain flexibility during R-loop initation. The presence of a shorter R-loop intermediate hints at an energetic barrier to forming subsequent base pairs, potentially acting as a bottleneck for target DNA binding. Incubation with the 12bp target produced a new intermediate with only 10 base pairs of the R-loop resolved (Fig. S3), correlating extension past the seed with high conformational flexibility, leading to a largely unresolved Rec2 domain.

The 16bp target continued to highlight the dynamic nature of the Rec2 domain and structural heterogeneity within the second half of the R-loop (Fig. S4). Within this dataset we captured a 15bp structure trapped in an unproductive state and a 16bp* structure (with 16 Watson Crick base pairs and a potential 17^th^ rC:dC base pair) with the NTS entering the RuvC active site. Both the 12bp and 16bp target datasets also had a small fraction of particles that yielded well resolved seed complexes (5bp and 8bp, respectively)—underscoring the significance of these early structures during initial DNA binding and reflecting the presence of additional structural barriers at intermediate R-loop lengths.

Finally, the 20bp target led to a structure of Cas12a with the full R-loop (Fig. 1c, Fig. S5). While this structure resembles previously published x-ray crystal structures^15,16,18^ and cryo-EM models^19^, because of the brief incubation time prior to vitrification, our structure is the first to capture the intact NTS poised for cleavage in the RuvC active site, elucidating activation of the nuclease domain. Together these structures highlight the dramatic conformational flexibility of Cas12a during R-loop propagation and DNA cleavage, consistent with previous fluorescence-based assays^20-24^ and molecular dynamics simulations^25^, and show how conformational changes are coordinated with R-loop propagation to create a highly specific enzyme.

### Cas12a uses a 5 base pair seed

The 5bp structure closely resembles the binary complex of Cas12a that contains five pre-ordered guide RNA nucleotides (LbCas12a: RMSD 5.3 Å)^17,26^, however, here the Rec1 domain has shifted up to accommodate formation of an early A-form heteroduplex (Fig. 2a). Within this 5bp structure Cas12a makes several non-specific contacts between the Rec lobe and the crRNA backbone (residues K15, R18, Y47, K51, N175, K530, F788, H872) (Fig. 2b), while the displaced nontarget strand is gripped by the PI domain. The TS and NTS rehybridize at the 6^th^ base pair positioned between the RuvC domain and the Rec1 helix-loop-helix, directing the DNA to exit the complex from the front of the protein. The DNA at the R-loop-DNA junction sits atop a bulky loop in the RuvC domain with K1054 projecting into the minor groove of the DNA (Fig. 2b). Interestingly, one structural intermediate from the same dataset contained a nearly identical structure but with a 5bp R-loop and an unresolved Rec1 domain (density is nearly absent starting at residue E49 where Rec1 would bend around the R-loop, -Fig. S2). This reconstruction marked by Rec1 flexibility likely represents Cas12a after initial PAM binding and R-loop initaiton and demonstrates the requirement for Rec1 flexibility during target search. R-loop initiation is then followed by Rec1 docking.

**Fig. 2.**
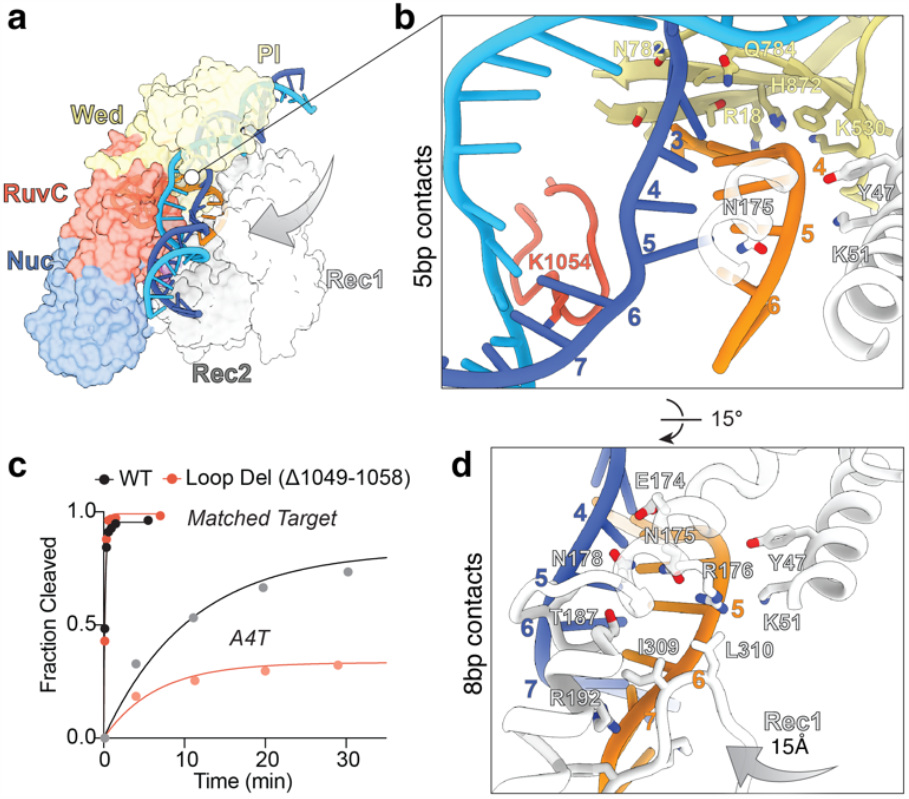
Cas12a binds to a 5bp seed. **a**, Structure of Cas12a with 5bp R-loop formed shown in surface representation. Arrow demonstrates the Rec1 docking movement. **b**, Enlarged view of the 5bp R-loop from panel **a** highlighting the Wed, Rec1, and RuvC contacts. Portions of Rec1 that do not contact the R-loop have been removed for clarity. **c**, WT and LD NTS cleavage of PT (matched, 20bp target) and a single mismatch target A4T. On-target cleavage is shown with dark data points, off-target cleavage shown with lighter data points. **d**, Enlarged view of the additional Rec1 R-loop contacts made in the 8bp structure. Arrow shows the inward rotation of the Rec1 domain.

The existence of this early 5bp intermediate despite additional target complementarity suggests that a structural barrier exists to limit R-loop propagation. The RuvC loop that bolsters the distal DNA could produce this barrier by acting as a bottleneck to allow for efficient DNA release if there is insufficient target complementarity. On the other hand, the RuvC loop could stabilize a transient early intermediate that precedes the energetic barrier to R-loop propagation, thus shifting the equilibrium of observed early R-loop intermediates to promote extension. To test the role of this RuvC loop, we purified a loop deletion mutant (LD, Δ1049-1058) and measured its effect on cleavage of a matched and mismatched target. LD did not have a measurable effect on NTS cleavage of the matched target (5.1 ±0.4 min^-1^ vs 6.4 ±1.2 min^-1^ for WT) but resulted in reduced cleavage of a single nucleotide mismatch (A4T) at position 4 of the R-loop (Fig. 2c, Table S3). The minor reduction in cleavage rate of the mismatch substrate reports that the RuvC loop was contributing to the stability of the 5bp intermediate rather than acting as a barrier to R-loop propagation. The increase in observed specificity by the LD mutant likely arises by destabilizing the early R-loop intermediate and increasing the rate of R-loop collapse for both DNA substrates, resulting in a measurable decrease in the presence of mismatches. This data shows that the RuvC loop is involved in increasing the efficiency of initial target search by stabilizing this early R-loop intermediate.

As the R-loop extends to 8bp, the Rec1 domain rotates toward the Nuc lobe by ∼15 Å and encircles the R-loop (Fig. 2d), forming a dense network of new contacts along the crRNA backbone from positions 5-8 (R176, R192, F306, K307, I309, L310) and stabilizing the TS (E174, N178, I185, S186). This early intermediate resembles previous Cas12a structures with an 8bp R-loop (Lb: RMSD 5.3 Å)^19,26^. R-loop extension past the initial seed coincides with Rec2 eviction: in the 8bp structure Rec2 has rotated outward ∼10 Å. This conformational change could contribute to the structural barrier that antagonizes extension past the 5bp seed. Interestingly, in both the 5bp and 8bp structures, density for the unpaired crRNA guide continues past the nascent R-loop, suggesting that there is a repeated pre-ordering mechanism to promote efficient R-loop propagation to subsequent bases.

For various RNA-guided nucleases, the first few nucleotides of the guide RNA (often exposed and pre-ordered) are noted as the “seed” due to their importance in preventing off-target binding during initial guide:target base pairing^27-30^. Here, our shortest R-loop intermediate matches the length of pre-ordered RNA in previous binary Cas12a structures^17,26^. We propose that the early 5bp intermediate represents a conformation with fast binding and dissociation kinetics used to rapidly check initial target complementarity upon PAM binding, acting as a seed for R-loop initiation^7^. The RuvC loop helps stabilize the 5bp seed before Rec1 docks and forms contacts with the extending R-loop. This short R-loop (∼8bp) intermediate is sufficiently stable to overcome the compact nature of Cas12a and promote dislodging of Rec2 and further R-loop propagation.

### Rec2 flexibility accommodates R-loop extension but delays contacts

Rec2 domain conformational dynamics is one of the most dramatic series of changes seen within the R-loop intermediate structures. In the 8bp structure, Rec2 rotates outwards to avoid steric clashes with the extending R-loop (Fig. 3a), and weak domain density indicates low occupancy due to flexibility. Rec2 then becomes completely unresolved in the 10bp structure, and the 16bp target leads to structures with Rec2 either unresolved, contacting the intermediate R-loop in an incorrect orientation, or weakly docking onto the R-loop (Fig. S4). An important consequence of the dynamic nature of Rec2 is the lack of protein contacts that are made in parallel with base pair formation. Only after the 16^th^ base pair is formed does Rec2 begin to dock onto the R-loop and make upstream contacts throughout the R-loop via the backbone and minor groove (Fig. 3b).

**Fig. 3.**
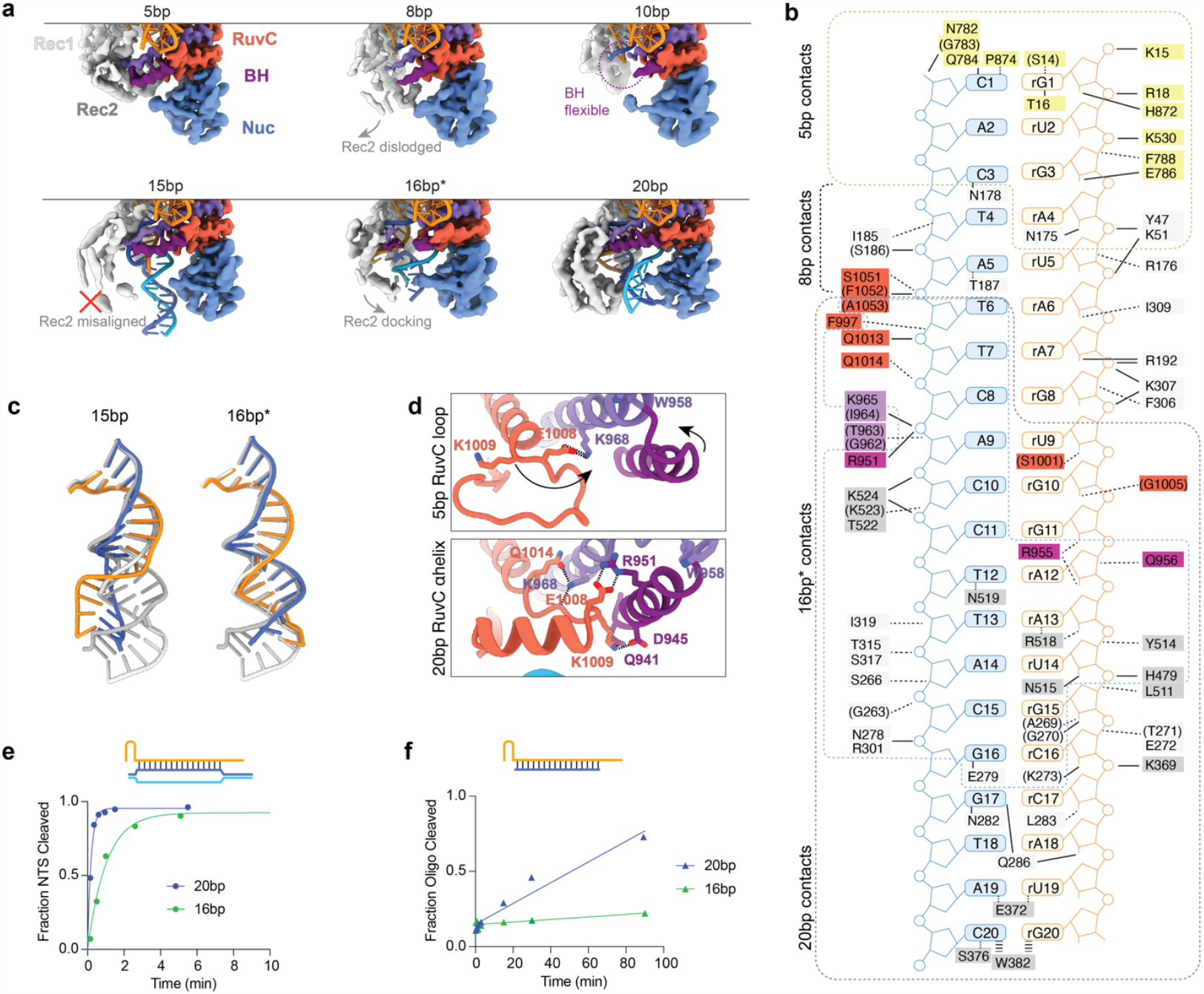
Middle R-loop propagation is marked by Rec2 flexibility, delaying R-loop contacts and RuvC activation. **a**, Reconstructions of Cas12a Rec and Nuc lobes during R-loop propagation. Protein is shown as reconstruction maps and nucleic acids are shown as model. WED and PI domains were removed from these images for clarity. Rec2 becomes dislodged as R-loop extends past the seed and returns to a cleavage-competent conformation after R-loop reaches 16bp. BH domain flexibility is also seen. **b**, Diagram of 20bp R-loop contacts grouped according to resolved length of the target (5bp: yellow, 8bp: black; 16bp*: blue; 20bp: grey). Amino acids are colored according to Cas12a domains as in Fig. 1. Black lines mark hydrogen bonds and dashed lines represent other contacts. Residues in parentheses denote backbone (not side chain) contacts. **c**, Comparison of late R-loop intermediates to the 20bp R-loop (grey). **d**, Enlarged side view of RuvC lid and BH highlighting the change in contacts made by E1008 and K1009. Arrows show the change in position of the BH and RuvC lid between the 5bp and 20bp models. **e**, Example time course of WT Cas12a NTS cleavage of DNA substrates that form a 16bp or 20bp R-loop. **f**, Example time course of WT Cas12a trans-cleavage when activated with single stranded oligos that form a 16bp or 20bp R-loop.

In addition to Rec domain conformational flexibility, the growing R-loop is centrally located within the Cas12a channel due to helical deformations that diverge from the path of the final 20bp R-loop heteroduplex (Fig. 3c, Fig. S4). This contrasts with the first half of the R-loop, which remains in an A-form helix. As captured in the 15bp structure, starting at position 10, the minor groove narrows and the heteroduplex stretches to accommodate R-loop propagation compatible with the constraints imposed by the continuous NTS lining the Nuc lobe. Thus, the nascent heteroduplex is incompatible with Rec2 docking before forming the last few R-loop base pairs. One structural consequence of Rec2 domain flexibility is the lack of rigidity observed for the BH domain (here considered inclusive of RuvC II helix 1). Our R-loop intermediate structures show that the BH starts to dislodge from Rec2 in the 8bp structure, resulting in conformational flexibility during middle R-loop propagation that allows the conformational transition seen in the final 20bp structure (Fig. 3a).

In our 16bp* structure, new protein contacts form along the R-loop via the phosphodiester backbones and the minor groove. Rec1 shifts inwards to the R-loop, forming new contacts (G263, S266, A269, G270, K273, N278, E279, R301, T315, S317, I319). Rec2 begins to dock (via H479, Y514, N515, R518, N519, T522, K523, K524), but insufficient contacts prevent stable docking into the final conformation. The BH, rearranged and anchored into Rec2 via W958, also begins to dock onto the middle R-loop with several contacts (R951, R955, Q956). The rearrangement of helices in the BH appears to occur before R-loop docking, as it is seen in the 15bp structure despite incorrect Rec2 docking preventing R-loop contacts. Finally, formation of the last R-loop base pairs is accompanied by additional contacts that favor stable docking (Rec1: T271, E272, N282, L283, Q286; Rec2: K369, E372, S376, W382 base stacking, L511; BH: G962, T963, I964).

The observed flexibility of Cas12a and the heterogenous nature of the middle R-loop intermediates shows propagation past the early R-loop proceeds without protein contacts guiding heteroduplex formation. Instead, R-loop propagation must be driven by the energetics of DNA base pair melting and RNA:DNA base pair formation.

### Late R-loop Rec2 docking promotes RuvC exposure

Domain docking onto the R-loop stabilizes the ternary complex and reorients the domains relative to one another, as seen in the 16bp* and 20bp R-loop structures. As Rec2 docks onto the R-loop, the BH anchors in an orientation that causes its two helices to shift upwards and rotate inwards towards the R-loop (Fig. 3d). This conformational rearrangement positions the BH to interact with the R-loop and the adjacent RuvC lid, stabilizing the newly observed *α*-helix conformation that exposes the RuvC active site. In early R-loop intermediates, lid K1009 points away from the BH, but when the RuvC lid transitions into an *α*-helix in the 20bp structure, K1009 contacts the BH via Q941 and D945. Additionally, lid E1008 is pulled up by BH R951 in the 20bp structure, supporting the raised conformation of the lid. Consistent with a model in which Rec2 docking contributes to RuvC active site exposure, unstable Rec2 docking in the 16bp* structure results in structural heterogeneity in the BH and RuvC lid, causing less-well resolved density for the RuvC lid (Fig. S7a). Further, the 15bp R-loop structure shows that incorrect Rec2 docking prevents the BH from making RuvC lid contacts and the lid remains as a loop blocking the active site.

Our structural observations are consistent with previous biochemical characterization of the BH’s (RuvC II helix 1) effect on ternary complex formation and RuvC activity^23,24^. According to our model, R-loops with sufficient lengths to permit semi-stable Rec2 docking will achieve RuvC activation but at reduced efficiencies caused by Rec2—and BH—flexibility. To test this, we performed cleavage reactions with reduced R-loop lengths. NTS cleavage of the 16bp target duplex (with the potential to form a 17^th^ rC:dC base pair) occurred at a 4-fold reduced rate compared to that of a 20bp R-loop (Fig. 3e, Table S3). Because rate-limiting R-loop formation of the 20bp target could mask penalties on cleavage, we also tested trans-cleavage by Cas12a activated with single stranded oligos to isolate the RuvC exposure step from rate-limiting R-loop formation. The 16bp R-loop resulted in 22-fold slower rates than the 20bp R-loop (8.1 ±1.1 ×10^5^ M^-1^ min^-1^) which is in agreement with the ≥4-fold penalty observed for NTS cleavage (Fig. 3f). Together, these data highlight the effect of R-loop length on RuvC activation via Rec2 docking.

Comparison to early R-loop intermediates shows how RuvC catalytic residue accessibility relies on the RuvC lid loop-to-*α*-helix transition. In the 20bp structure, lid K1000 orients away from the active site and interacts with RuvC Q1013 and E1016 to further reinforce the RuvC lid *α*-helix conformation. In earlier intermediates, K1000 is responsible for interacting with and sequestering catalytic residue E993 from the active site (Fig. S7b). In early R-loop intermediates, D1263 is drawn away from the active site by R1226 and N913. As the RuvC lid transitions into an *α*-helix, R1226 moves to interact with the incoming DNA and N913 is pulled away from D1263 so that the catalytic residue can rotate towardsthe active site.

Of note, both our kinetics-guided R-loop structures with the RuvC active site exposed also have the NTS in the active site. These structures (and the lack of an exposed, unoccupied RuvC active site) hint at a role of the NTS in actively displacing the unstructured RuvC lid. In support of this hypothesis, the 16bp* lid is displaced and the NTS is in the active site before RuvC lid-BH contacts have formed. In this case, Rec2 docking stabilizes the displaced RuvC lid in an *α*-helix conformation that leads to correct RuvC activation and substrate positioning for efficient cleavage (discussed below).

We propose Rec2 docking late in R-loop formation represents a conformational checkpoint leading to Cas12a activation. Our structures show stable docking occurs once the R-loop has reached a sufficient length and correct geometry. Off-target DNA sequences could introduce R-loop helix distortions, such as kinks or minor groove changes, limiting the stability of Rec2 in the docked state^31,32^. This late conformational checkpoint establishes a cleavage-competent conformation of Cas12a, enabling specificity against mismatches late in the R-loop.

### Nontarget strand poised for cleavage in the RuvC active site via base stacking interaction

In addition to stable protein docking, R-loop completion also causes the distal target DNA to move from the center of Cas12a to exiting through the back. This directional movement of the DNA and simultaneous increase in single stranded NTS length causes the NTS to traverse across the RuvC domain during R-loop completion and displace the RuvC lid that had originally occluded the RuvC active site (Fig. 4a). Kinetically relevant sample preparation enabled capturing a 20bp structure with the NTS in the RuvC active site poised for cleavage.

**Fig. 4.**
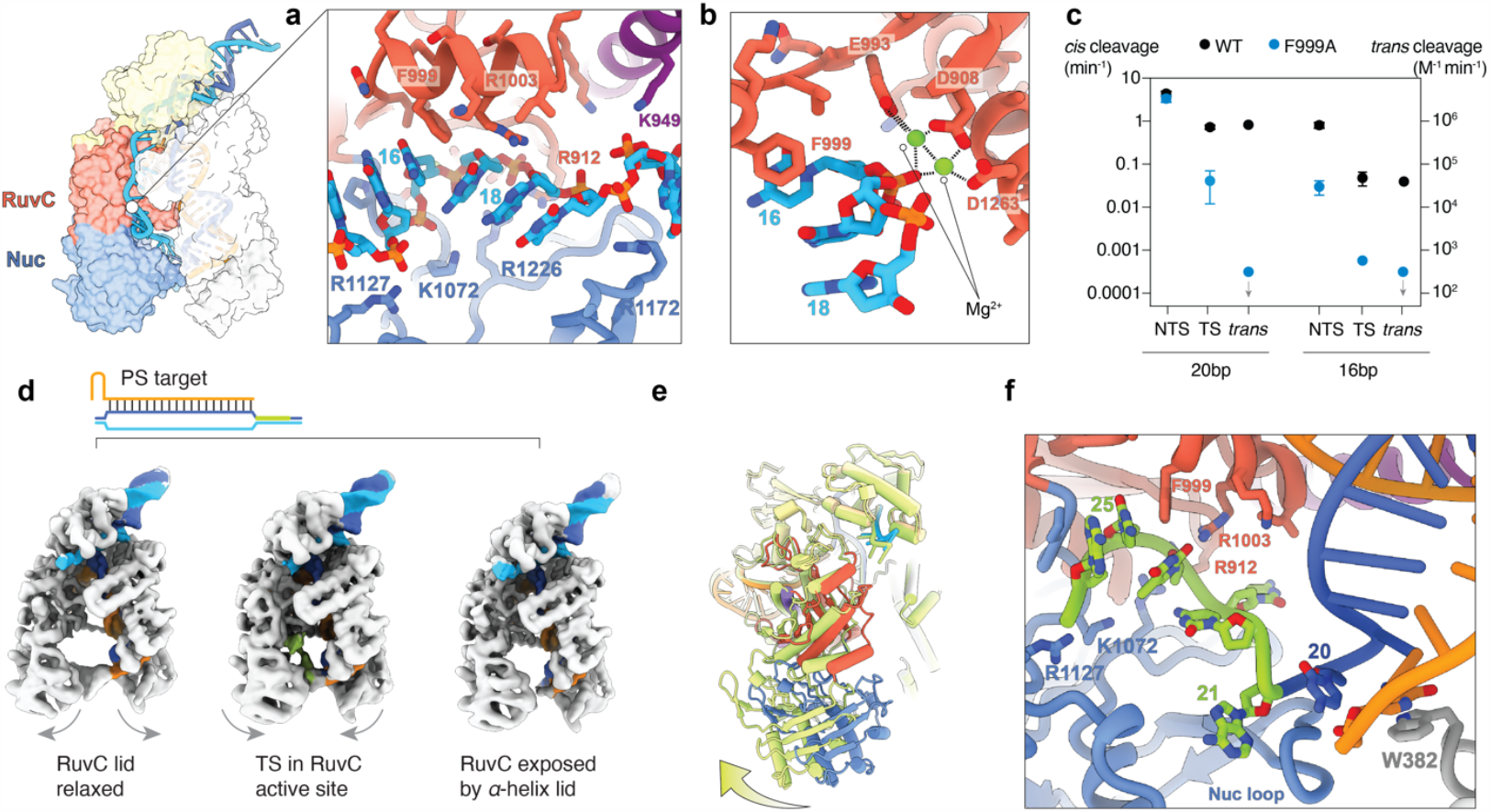
DNA cleavage by Cas12a relies on F999 base stacking. **a**, Enlarged view of the model for the 20bp structure showing the NTS in the RuvC active site. A network of positively charged residues surrounding the NTS bring the substrate to the active site. F999 base stacks with base 16 of the NTS upstream of the scissile phosphate, kinking the scissile phosphate into the active site. **b**, RuvC active site is poised for NTS cleavage as the three catalytic residues (D908, E993, D1263) coordinated with two Mg^2+^ ions and the scissile phosphate. **c**, Cleavage inhibition by F999A mutant. Observed cleavage rates for NTS and TS are plotted on the left y-axis and second order rate constants for trans-cleavage are plotted on the right y-axis. Data points with a gray arrow represent upper limits to the rate as no cleavage was observed. **d**, EM maps reconstructed from a target DNA with a phosphorothioate-modified TS backbone (green). Structures show Cas12a with the TS in the active site (right), Cas12a with the active site exposed and no TS (left), and Cas12a in an extended conformation in which the RuvC lid is not observed as a stable *α*-helix. **e**, Cas12a with TS in the active site (colored as in Fig. 1) overlayed with Cas12a in an extended conformation (green). Models were aligned along the R-loop and Rec domains to show movement in the Nuc lobe, particularly the Nuc domain. **f**, Enlarged view of the TS in the RuvC active site. TS base 20 is no longer base paired in the R-loop and base 25 is base stacking with the lid F999. Protein is shown in cartoon and surface representation to demonstrate interaction pocket.

The NTS is positioned upstream between the Rec1 helix-loop-helix and RuvC domain, propped up downstream by the Nuc loop, and secured across the RuvC active site by a network of positively charged residues (R912, K949, P1068, K1072, R1127, R1172, R1226, N1295). The narrow pocket between the RuvC and Nuc domains enforces a bent conformation in the DNA. The NTS engages the RuvC lid *α*-helix solely via F999 and R1003 (Fig. 4a, Fig. S8a). F999 base stacks with the NTS upstream of the scissile phosphate, modeled as position 16, stabilizing the kinked DNA in the active site. R1003 contacts the bases downstream from the scissile phosphate, responsible for both attracting the NTS and stabilizing the DNA shape. The RuvC catalytic residues D908, E993, and D1263 coordinate with the scissile phosphate using two Mg^2+^ ions (observed as continuous density) (Fig. 4b, Fig. S8b). This catalytic architecture resembles what has been seen for other class 2 RuvC nucleases coordinated for cleavage^33,34^.

To understand the role of the RuvC lid, we tested Cas12a mutants that disrupt lid-DNA contacts seen in the 20bp structure. We made an alanine substitution of F999 and tested RuvC-mediated cleavage. Surprisingly, there was no defect observed for NTS cleavage of the 20bp target by the F999A mutant (Fig. 4c, Fig. S8d) and we reasoned that a penalty could be masked by other mechanistic features (rate-limiting R-loop formation, network of positive residues). To test this hypothesis, we repeated the cleavage assay with the 16bp target. NTS cleavage of the 16bp target showed a substantial 25-fold defect for the F999A mutant compared to WT Cas12a (Table S3). TS cleavage by F999A showed a 9-fold defect for the 20bp target and 130-fold defect for the 16bp target. These cleavage data, combined with our structures, demonstrate the importance of the lid phenylalanine base stacking interaction for stabilizing the kinked DNA to ensure efficient cleavage. To observe penalties more directly on RuvC activation, we compared trans-cleavage by F999A complexed with single stranded activators. In stark contrast to WT Cas12a, the F999A mutant showed no cleavage of the trans substrate after 48 hours when activated with a 20bp or 16bp R-loop (Fig. 4c). This complete loss of measurable activity underscores the importance of F999 base stacking in stabilizing the DNA substrate within the RuvC active site.

We also tested removal of the second lid-NTS contact with R1003A (Fig. S8d). Cleavage penalties were not observed when Cas12a R1003A was activated with a 20bp R-loop. However, with a 16bp R-loop, R1003A cleaved the NTS and the TS 15-fold and 7-fold slower than WT, respectively. Validating structural observations, these cleavage data confirm that both lid contacts play an important role in RuvC-mediated DNA cleavage.

### Cas12a dynamics allow target strand positioning in RuvC

An outstanding question in Cas12a DNA targeting is how the TS reaches the distant RuvC nuclease for double stranded DNA cleavage. To achieve the same polarity as the NTS and access the RuvC active site, the DNA downstream of the R-loop must melt several base pairs and the TS must sharply bend. Molecular dynamics simulations suggest that the RuvC lid *α*-helix anchors the middle R-loop while the Rec2 and Nuc domains use coordinated movements to guide the TS into the active site^35^. To provide direct structural insights into this process, we prepared Cas12a with a 20bp target modified with phosphorothioate (PS) linkages at potential TS cleavage sites to prevent cleavage and product release. Samples were incubated for 30 minutes (sufficient to reach 100% cleavage of unmodified target DNA) before vitrification. From this sample, we obtained three distinct structures with nominal resolutions ranging from 3.5-3.6 Å (Fig. 4, Fig. S9-10, Table S1).

Post-NTS cleavage states are characterized by dramatic Rec2 flexibility. Within the PS dataset, a substantial class of particles resulted in a structure aligned with a fully formed R-loop and cleaved NTS, but without a resolved Rec2 domain (Fig. S9). Three-dimensional variability analysis produced reconstructions with continuous density between the downstream nucleic acid and the Nuc domain loop, suggesting the Nuc domain remains bound to the distal DNA. Consistent with this structural class, recent single molecule experiments characterized a Nuc-driven clamped state that co-occurs with NTS (and TS) cleavage before DNA release and suggested Rec2 dissociation from the R-loop following cleavage^36^.

Of the reconstructions with a Rec2 domain docked, three-dimensional classification led to three distinct classes of Cas12a: (1) an extended conformation with the RuvC lid relaxed; (2) a closed conformation with the TS captured in the RuvC active site; (3) and a closed conformation with the RuvC active site exposed and no bound nucleic acid (Fig. 4d, Fig. S10). The ‘closed conformation’ structures closely resemble the 20bp R-loop reconstruction with the NTS in the active site (RMSD 0.71 Å and 0.57 Å). In the extended structure the two lobes move outward, with the Nuc domain rotating towards the back by ∼9 Å, and the Rec2 domain moving ∼5 Å (Fig. 4e). Here, the R-loop moves with the Rec lobe but remains in contact with the Nuc loop.

We observe contiguous density for the TS from the R-loop to the RuvC domain (Fig. 4f, Fig. S8). The TS, single stranded and in the right orientation, is brought to the RuvC domain by a small network of positively charged amino acids in the Nuc domain (P1068, R1127, R1226). The RuvC lid R1003 and F999 interact with TS similarly to the NTS—stabilizing the scissile phosphate in the active site. This structure agrees with our cleavage data and underscores the importance of the lid in securing the TS for cleavage. Here, the catalytic residues are not resolved in their activated state. Interestingly, we see the terminal R-loop base pair melts to enable TS bending, as predicted by MD simulations^35^. The structure with the RuvC active site exposed and no nucleic acid bound likely represents a transient state after DNA dissociation. Consistent with a previous hypothesized conformational state^36^, the Nuc domain loop extends to the base of the R-loop making substantial contacts to maintain a closed conformation.

Three-dimensional variability analysis of the consensus ‘extended’ reconstruction demonstrated the movement of density representative of the downstream nucleic acid bending towards the front of the protein, bringing the TS closer to the RuvC domain (Fig. S9). Here, the RuvC lid is unresolved due to the lack of contacts from the BH and nucleic acids, but the residues flanking the lid downstream are in the conformation associated with the unstructured loop that occludes the RuvC active site (and at low signal contour thresholds can be traced as the unstructured loop). This observation suggests that the RuvC active site does not remain exposed following initial cleavage, resetting between each subsequent cleavage event. Consequently, subsequent RuvC activation would be dependent on the repeated docking of the Rec2 (and BH) domain onto the R-loop. A RuvC active site that does not remain exposed for subsequent cleavage events could explain the greater penalties measured for TS vs NTS cleavage by F999A and R1003A.

Due to blocked TS cleavage, these structures represent transient conformational states that exist along the reaction pathway between NTS and TS cleavage. While Rec2 docking is important for exposing the RuvC active site to both DNA strands, Rec2 dynamics are needed for remodeling the downstream DNA. Our structures support the proposed coordinated movements of the Rec2 and Nuc domains to lift the DNA, creating a bend in the TS that is needed for reaching the RuvC domain^35^.

## Discussion

Here, we have captured a series of Cas12a R-loop structures that represent R-loop propagation on pathway to DNA cleavage. Kinetics-guided sample preparation allowed us to capture two unique structures with the NTS in the RuvC active site. We then used a chemically modified target to block cleavage of the TS and observed conformations on pathway to TS cleavage. Together, our structures represent a near complete reaction pathway, from R-loop initiation to NTS cleavage and subsequent TS entry into RuvC (Fig. 5).

**Fig. 5.**
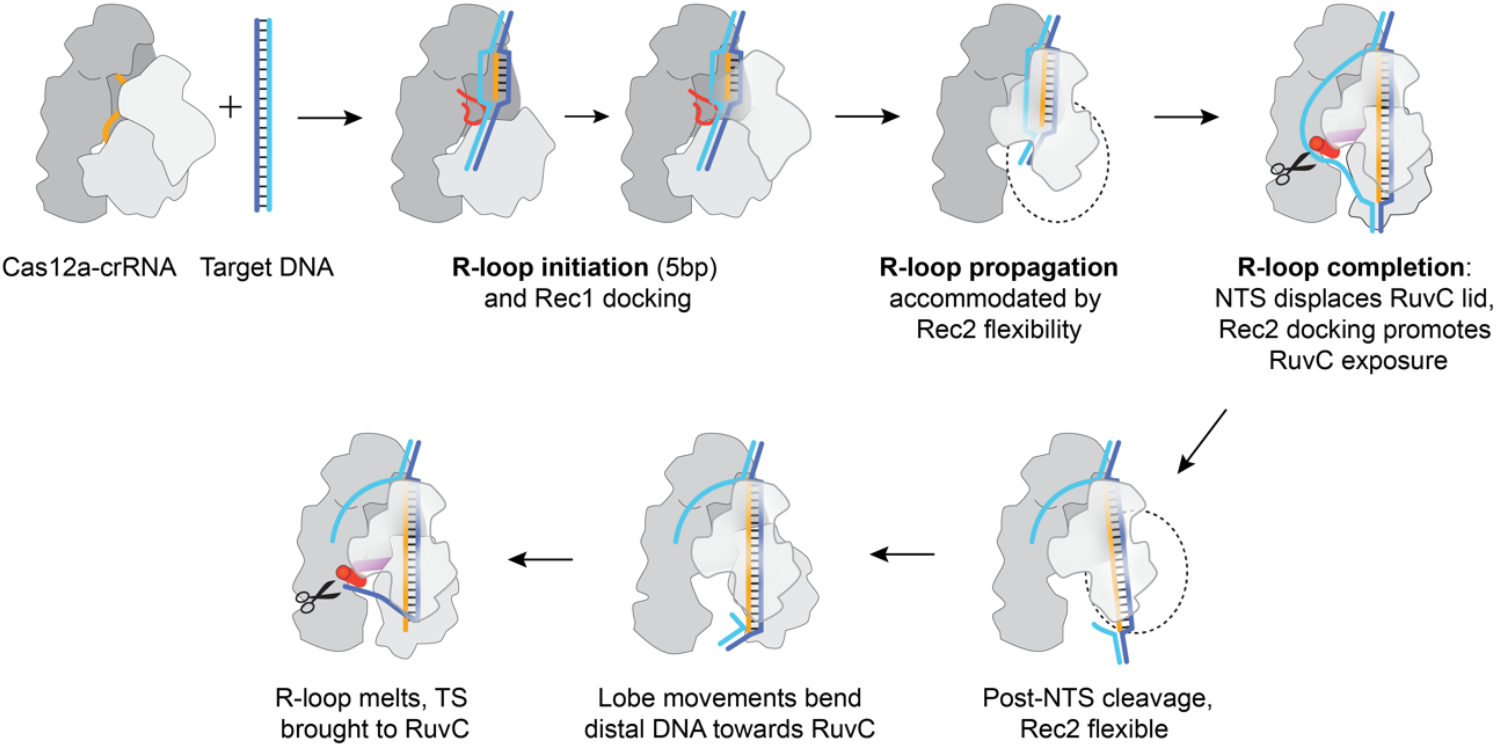
Structural model for Cas12a R-loop formation and DNA cleavage. Upon PAM recognition, Cas12a interrogates the first 5 bases of target DNA to search for seed complementarity. Protein contacts via Rec1 and the RuvC loop stabilize the seed to promote R-loop extension. Rec2 dislodges to accommodate R-loop propagation and remains flexible until the R-loop reaches sufficient length (∼16 to 20bp) to enable Rec2 (and BH) docking. R-loop completion brings the NTS across the RuvC domain, displacing the RuvC lid and exposing the active site. An alpha-helix RuvC lid uses F999 base stacking to help position and stabilize the scissile phosphate. NTS cleavage leads to Rec2 flexibility which—in coordination with the Nuc domain—brings the TS to the RuvC active site.

Our structures of early R-loop formation provide insight into the Rec dynamics of R-loop initiation and capture a 5bp seed. This 5bp seed corresponds to large mismatch penalties on Cas12a binding kinetics and free energy, however large penalties are also seen beyond the first 5bp of the R-loop ^7^. This discrepancy requires Cas12a to use other strategies to ensure specific target binding and cleavage. Interestingly, the early Cas12a R-loop intermediates are reminiscent of the recently observed SpCas9 bipartite seed: removal of the protein structures that appeared to create a barrier to early R-loop propagation actually increased specificity, contrary to expectations for removal of a structural barrier^37^. This comparison is another example of how class 2 nucleases have evolved similar yet distinct mechanisms to ensure efficient DNA targeting.

Subsequent extension through the middle of the R-loop is accommodated by dramatic Rec2 conformational flexibility, which leads to minimal R-loop contacts from both Rec2 and the BH. Previous analysis of mismatches within the Cas12a R-loop calculated large energetic penalties on equilibrium binding through most of the R-loop, within range of penalties expected by Nearest Neighbor rules for a duplex in solution^7,38,39^. This similarity in behavior suggested that the R-loop is being formed as it would in solution, with no protein contacts guiding or interfering with the energetics of base pair formation. Our structures help rationalize this model for describing propagation through the middle of the R-loop.

As the R-loop of Cas12a forms the final few base pairs, Rec2 begins to dock onto the R-loop. Domian docking provides numerous contacts throughout the R-loop and stabilize the cleavage-competent conformation. The delayed occurrence of these contacts relative to base pair formation establishes R-loop formation and RuvC activation into discrete steps along the reaction pathway and represents a conformational checkpoint for Cas12a activation. These contacts include the “linker” and “finger” checkpoints previously described for FnCas12a^19^. Again, we draw a mechanistic parallel for extending specificity to late in the R-loop analogous to Cas9^32,40,41^.

Considering recent Cas9 R-loop studies^31,32,37^, Cas12a appears to deviate in R-loop formation strategies, which likely contributes to distinct specificity profiles. Structures of Cas9 R-loop intermediates suggest Cas9 contacts R-loop base pairs nearly as they form. Following R-loop seed formation, Cas12a delays additional R-loop contacts until near R-loop completion. Consistent with these structural insights, Cas12a binds more specifically than Cas9, but suffers from reduced rates of R-loop formation and cleavage^7,8,11-13^.

Following R-loop formation, we were able to capture both the NTS and the TS in the RuvC active site of WT Cas12a. Both structures were highly similar to one another and relied on Rec2 docking to the R-loop. Both show the base upstream of the scissile phosphate base stacking with F999 of the RuvC lid, similar to previously published class 2 RuvC nucleases^32-34,42,43^. For NTS cleavage, our two structures suggest that both BH contacts and the NTS substrate itself contribute to promoting the RuvC lid loop-to-*α*-helix transition to expose the active site. For TS cleavage, Rec2 motions work with the Nuc domain to promote bending and lifting the TS into the RuvC active site.

Interestingly, the Rec2 dynamics seen post-NTS cleavage were associated with flexibility in the RuvC lid: reconstructions of the extended or open conformations had poor or ambiguous density representing an unstructured loop, suggesting the RuvC active site does not remain exposed between cleavage events and must be re-engaged for subsequent cleavage. The dependence on BH-lid contacts and DNA substrate to stably expose the RuvC active site can explain why previous ternary structures of Cas12a resulted in various lid conformations^15-19^.

The insight from these intermediate structures contributes to a more complete mechanistic understanding of Cas12a DNA targeting. We uncovered dramatic dynamics and conformational checkpoints during R-loop formation and captured RuvC activation. The distinction between R-loop formation and cleavage activation could guide future efforts to tune Cas12a efficiency and specificity. These kinetically relevant structures of R-loop intermediates on pathway to DNA cleavage have given temporal resolution to the transient interactions and unprecedented dynamic nature of Cas12a DNA targeting.

## Methods

### Cas12a cloning and purification

Cas12a was cloned into a pET-based expression vector^8^ with an N-terminal 6xHis-MBP tag, lac-inducible promoter, and Kanamycin resistance. The His-MBP-Cas12a plasmid was transformed into BL21(DE3) cells (New England Biolabs). A single colony was used to inoculate LB media supplemented with 50μg/ml Kanamycin for an overnight culture grown at 37°C, 200 rpm. The starter culture was then passaged (100X dilution) to 2L of LB supplemented with antibiotic and grown to an OD_600_ of ∼0.6 at which point 1mM IPTG was added to induce expression at 18°C. Cultures were grown for an additional 20 hours. Cells were pelleted and lysed in equilibration buffer (1M NaCl, 20 mM HEPES, pH 7.5, 0.5mM TCEP, 5% glycerol) supplemented with 200mM PMSF, 0.1% Tween-20, c0mplete Protease inhibitor cocktail tablet (Roche). Lysate was then incubated with 10 mM MgCl_2_ and DNase at 4°C with constant shaking for 20 minutes. Lysate was sonicated, clarified by centrifugation at 18k rpm for 30 minutes at 4°C. Clarified lysate was applied to a HisTrap HP column (Cytiva). His-tagged Cas12a was washed with 10% elution buffer (1M NaCl, 20 mM HEPES, pH 7.5, 5% glycerol, 250mM imidazole) before eluted with a linear gradient elution. Pooled fractions were digested by recombinant TEV protease (purified in house) to remove the N-terminal His-MBP tag and dialyzed overnight into low salt buffer (150 mM NaCl, 20 mM HEPES, pH 7, 0.5mM TCEP, 5% glycerol) at 4°C. Cas12a sample was then run through a HiTrap SP HP column (Cytiva) and eluted by linear gradient high salt elution (1M NaCl, 20 mM HEPES, pH 7, 0.5mM TCEP, 10% glycerol). Cas12a was then fractionated over a S200 Increase 10/300 GL column (Cytiva) equilibrated with low salt buffer supplemented with 5mM MgCl_2_. After each chromatography step, fractions containing Cas12a were confirmed by SDS-PAGE and pooled. Samples were concentrated to ∼10uM and aliquots were flash frozen in liquid nitrogen and stored at -80C.

Cas12a mutants were cloned using Q5 polymerase and KLD kit (New England Biolabs) and purified the same way as WT AsCas12a.

### Target DNA substrates

DNA oligos were purchased from IDT and resuspended in TE. Target duplexes were formed using 1:1.2 TS:NTS in 50 mM HEPES, 100 mM NaCl and heating for two minutes at 90C and slow cooling to 25°C. When using 5’-FAM-labeled oligos, the complementary oligo was added in 1.2-fold excess and labeled duplexes were annealed the same way. DNA targets with reduced complementarity to the guide had PAM-distal mismatches introduced by inverting the TS:NTS base pair so that Watson Crick crRNA:TS base pairs could not form at these locations.

### Cryo-EM sample preparation, data collection

50μM crRNA (Synthego) was added to an aliquot of purified 12μM WT Cas12a at a ratio of 1:3μl and incubated at room temperature for 30 min. Equal volumes of 10μM duplex DNA and assembled Cas12a-crRNA were mixed and incubated at 37ºC or ambient temperature (∼18°C) before vitrification. DNA binding reaction incubation times varied depending on the DNA substrate used: 8bp and 12bp DNA substrates were incubated for 1 hour at 37ºC, 16bp DNA incubated for ∼4 minutes, and 20bp DNA incubated for 1 minute. 1.2/1.3R 400 mesh Cu grids were plasma-cleaned for 30 sec in a Solarus 950 plasma cleaner (Gatan). All cryo-EM samples (2.5ul) were applied to grids using an FEI Vitrobot MarkIV (Thermo Fisher) set to 4C and 100% humidity. Samples were blotted for 6 seconds at 0 force before being plunge frozen into liquid ethane and stored in liquid nitrogen.

The 20bp dataset was collected on a FEI Glacios cryo-EM microscope (200kV) equipped with a Falcon 4 direct electron detector (Gatan). Movies were recorded in SerialEM^44^ with a pixel size of 0.94Å and a total exposure time of 15 sec resulting in an accumulated dosage and 49e^-^/Å^2^ split into 60 frames. The 8bp, 12bp, and 16bp datasets were collected on a FEI Titan Krios (300kV) equipped with a K3 Summit direct electron detector (Gatan). Movies were recorded with SerialEM with a pixel size of 0.8332Å and a total exposure time of 3.8 sec resulting in an accumulated dosage of ∼80e^-^/Å^2^ split into 100 frames. All datasets were collected at 30° tilt and uploaded to cryoSPARC Live for initial real time processing. For the 8, 12, 16, 20bp datasets, 5,803, 3,492, 12,100, and 2,403 movies were collected, respectively.

### Cryo-EM data processing

All datasets were initially processed using MotionCor2^45^ and downstream processing was done in cryoSPARC^46^. Processing workflows are shown in Fig. S2-5, 10.

### Model building and refinement

Previously published AsCas12a was rigid body fit into the 20bp structure within ChimeraX^47^. Individual domains were then rigid body fit into structures with rearranged protein. Modeling was performed through iterative rounds of using Isolde^48^ and Coot^49^. Models were then subjected to real space refinement within Phenix^50^ for final modifications and a validation score.

### Cleavage time courses

Purified Cas12a and mutants were assembled with cognate crRNA (Synthego) in excess at room temp in assembly buffer (150mM NaCl, 50mM HEPES, pH 7, 5 mM MgCl_2_, 2mM DTT) for 30 minutes. Before cleavage reactions, DNA and Cas12a were pre-warmed to 37 °C. To start the cis-cleavage (NTS, TS) reaction, 5’-FAM-labeled duplexes in reaction buffer were combined with Cas12a-crRNA at final concentrations of 50nM active enzyme and 10nM duplex DNA and the reaction was carried out in a 37°C water bath. Reaction buffer used had the same composition as assembly buffer supplemented with 0.2 mg/ml molecular biology grade BSA to prevent signal loss due to the preferential sticking of product to tubes. At various time points, 2μl were sampled from the reaction and quenched in 4μl of 0.1M EDTA. Time points were resolved via capillary electrophoresis using an Applied Biosystems DNA sequencer (ABI 3130xl). Traces corresponding to substrate and product were analyzed to plot fraction cleaved over time. Cleavage time courses were analyzed using a single exponential curve fit (non-linear regression) on GraphPad Prism. Rates are reported per minute.

Activated Cas12a for trans-cleavage reactions was assembled as described above. Single stranded activator oligos were incubated with Cas12a-crRNA for 30 minutes to ensure complete binding before addition of radiolabeled trans substrate 5’-TTATT initiated the reaction (Table S2). Trans-cleavage reactions were initially tested with varying amounts of excess substrate over 10nM activated WT Cas12a. Replicates were performed at 10nM Cas12a and 50 nM single stranded trans substrate. At various time points, 2μl were sampled from the reaction and quenched in 4 μl of denaturing quench (60% formamide, 20 mM EDTA, 0.01% bromophenol blue, and 0.01% xylene cyanol). Quenched samples were resolved on a 20% denaturing PAGE gel. Substrate and product intensities were quantitated using a phosphorimager. Cleavage time courses were analyzed using a linear fit (non-linear regression) in Kaleidograph. Second order rate constants were calculated by dividing the observed cleavage rate by the Cas12a concentration. Additional control experiments (not shown) established that 50 nM substrate was subsaturating. For F999A trans-cleavage reactions in which no cleavage was observed, upper limit rates were calculated from an approximate detection limit of 1% cleavage at a 48hr time point. Reactions were repeated with a second trans substrate (C_10_) to validate results.

## Data Availability

Structures of the 5bp, 8bp, 10bp, 15bp, 16bp*, 20bp R-loop intermediates have been deposited in the EMDB with accession codes EMD-40441, EMD-40442, EMD-40443, EMD-40444, EMD-40445, EMD-40446, respectively. Associated atomic coordinates are deposited in the PDB with accession codes 8SFH, 8SFI, 8SFJ, 8SFL, 8SFN, 8SFO, respectively. Structures from the TS(PS) dataset have been deposited in the EMDB EMD-40447, EMD-40448, EMD-40449. Aassociated atomic coordinates are deposited in the PDB with accession codes 8SFP, 8SFQ, 8SFR.

## Acknowledgements

We thank Dr. Axel Brilot and Dr. Evan Schwartz at the Sauer Lab at UT Austin for cryo-EM assistance, Prof. Ken Johnson and Dr. Tyler Dangerfield for use of the Applied Biosystems DNA sequencer, Prof. Giulia Palermo and her lab members and members of the Taylor Lab for valuable discussion around the project. This work was supported by the National Institutes of Health R35GM138348 to D.W.T. and R35 GM131777 to R.R. and Welch Foundation Research Grant F-1938 grant to D.W.T.

## Author Contributions

I.S. conceptualized the project, prepared samples for and performed cryo-EM, processed and interpreted all structural data, and performed biochemical assays. C.M. helped purify and characterize Cas12a mutants. A.-H.N. performed and analyzed trans-cleavage assays. I.S., C.M., A.-H.N., R.R., and D.W.T. interpreted the data. I.S. and D.W.T. wrote the manuscript with feedback from other authors. D.W.T. secured funding for the project.

