## Supplemental Information for "Structural basis of Cas12a R-loop propagation on pathway to DNA cleavage"

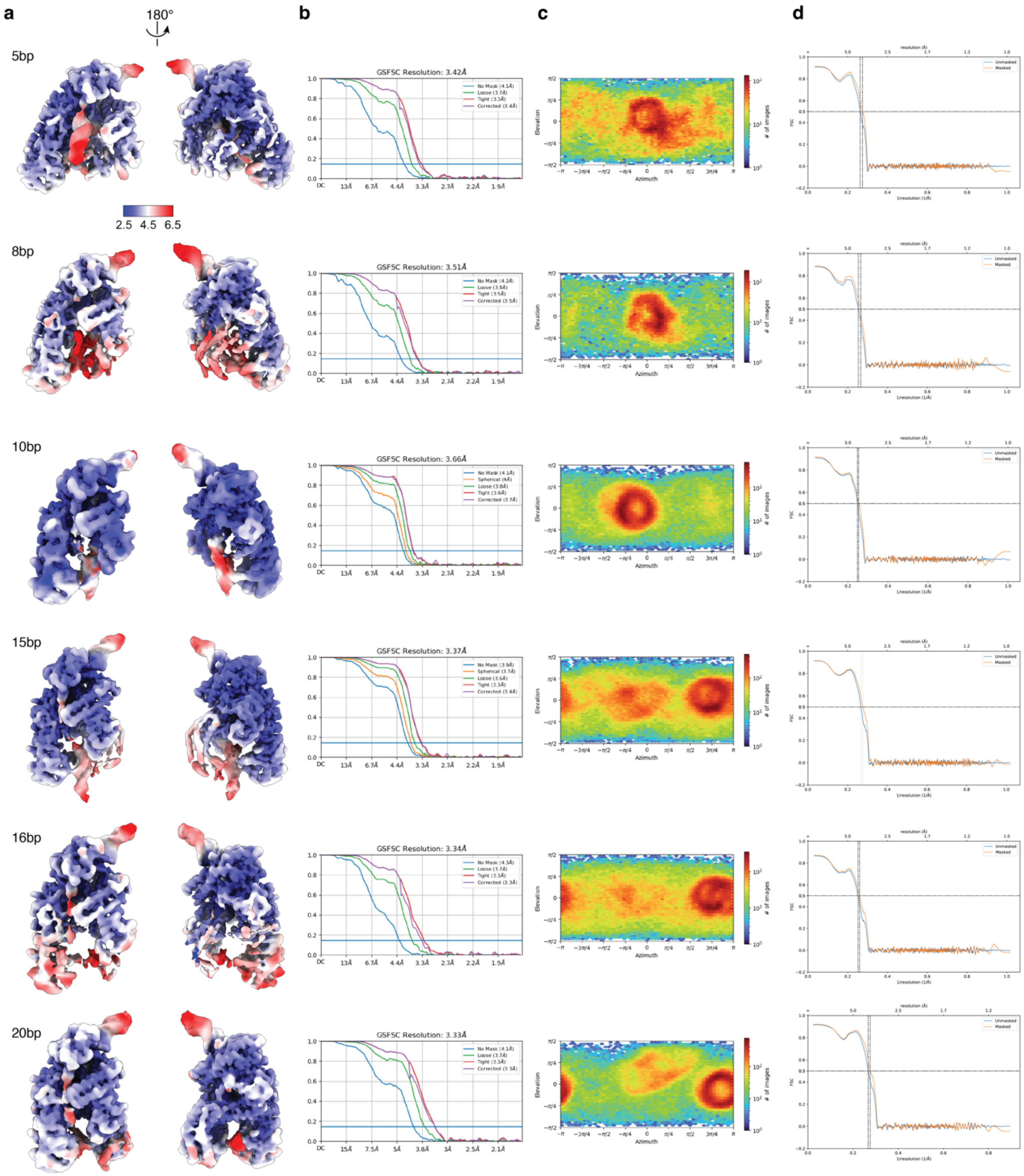


**Supp. Fig. 1 | Local resolution of intermediate R-loop maps. a,** Unsharpened maps of R-loop intermediates colored by local resolution. **b,** Gold-standard FSC curves for each reconstruction. Resolution estimated at FSC=0.143. **c,** Euler diagrams for each reconstruction showing particle orientation distributions. Map resolutions and particle distribution plots were produced in cryoSPARC. **d,** Map-to-model FSC curves produced in Phenix.

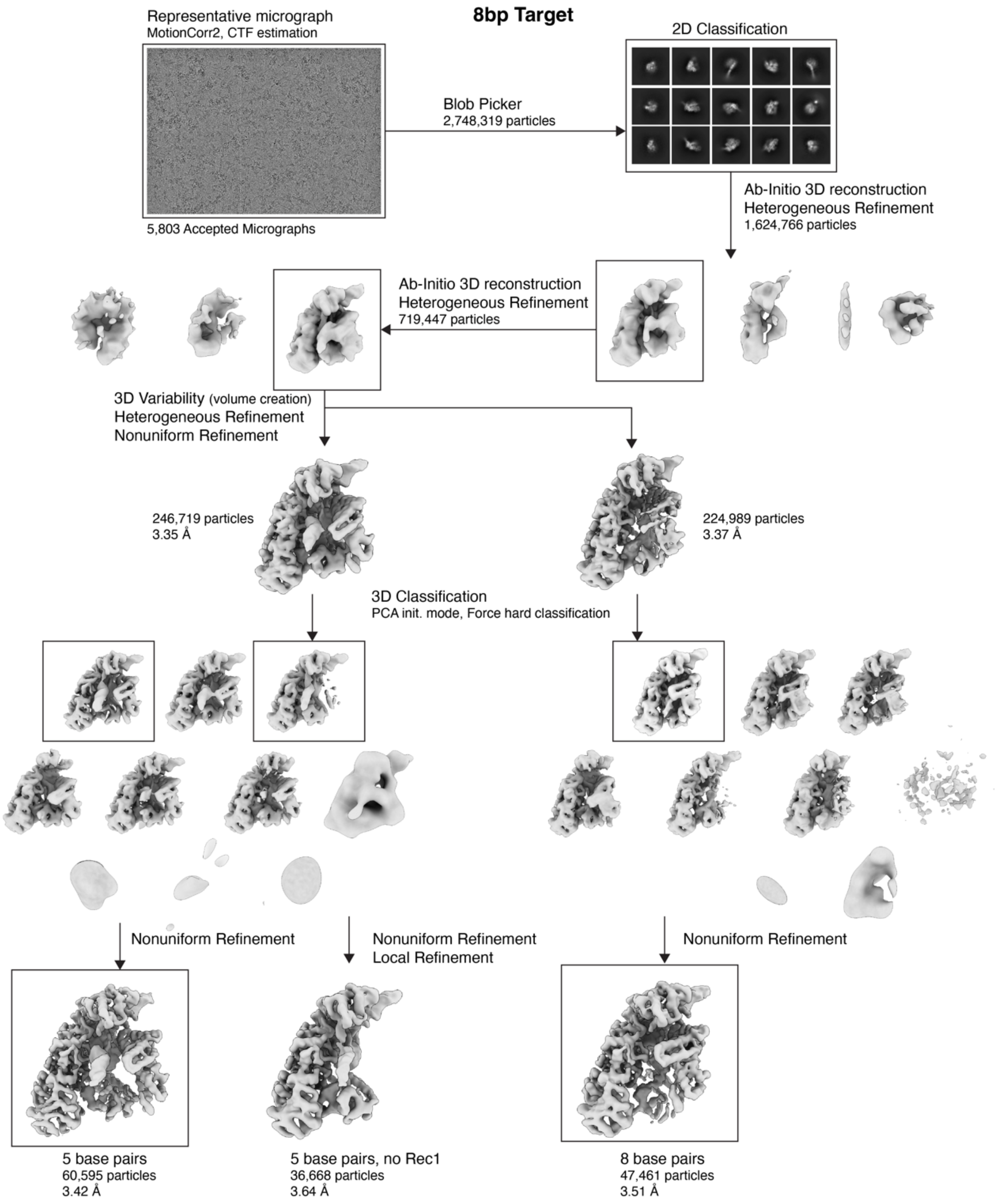


**Supp. Fig. 2 | Final data processing pipeline for 8bp target dataset.**

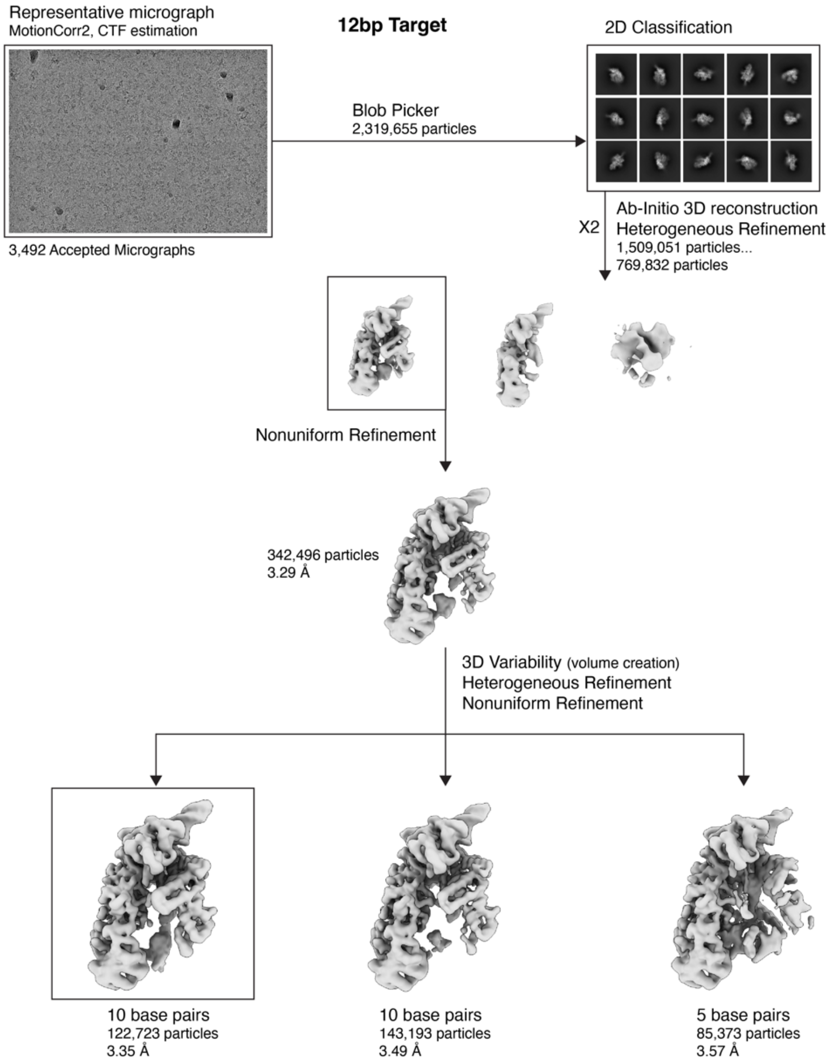


**Supp. Fig. 3 | Final data processing pipeline for 12bp target dataset.**

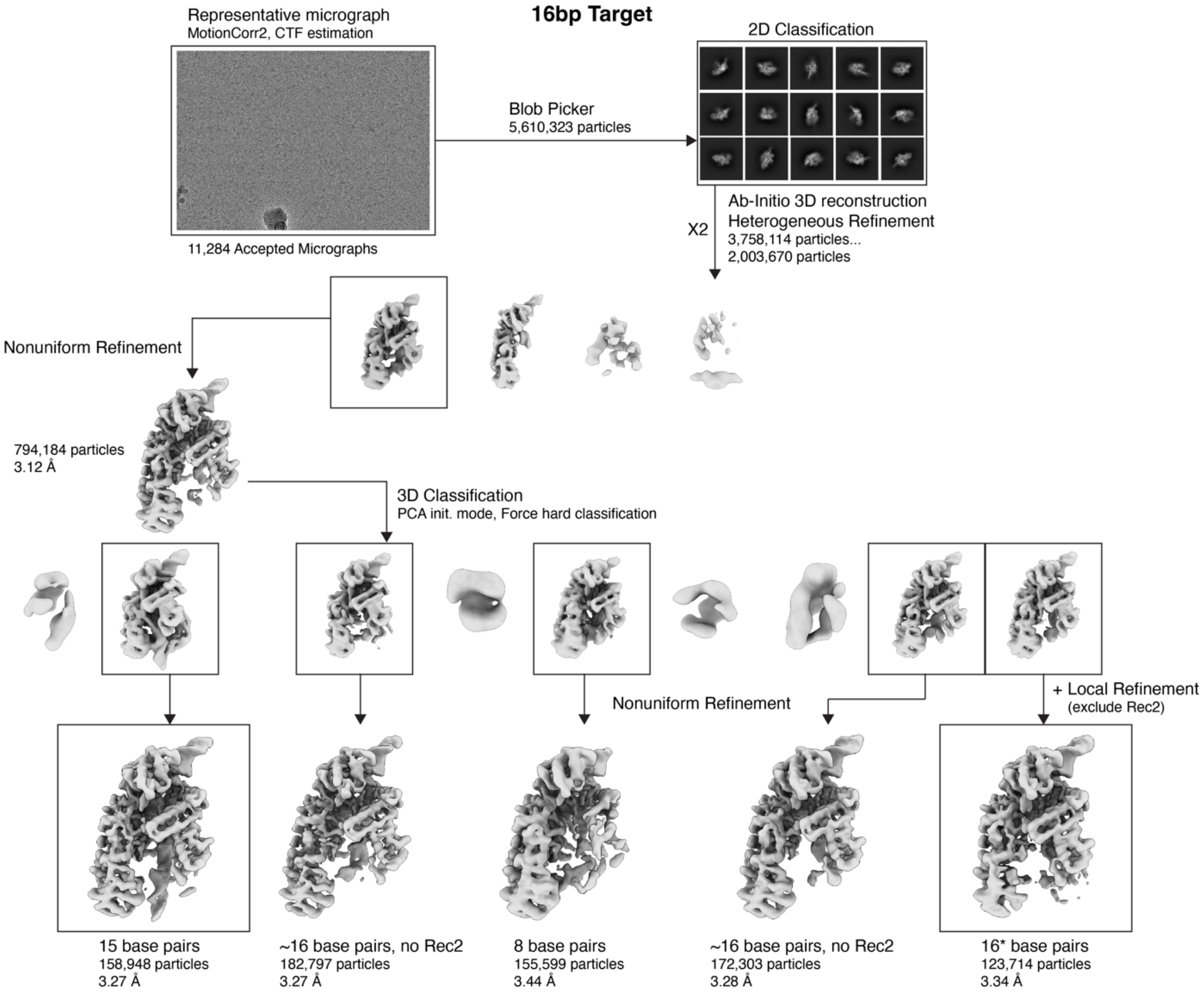


**Supp. Fig. 4 | Final data processing pipeline for 16bp target dataset.**

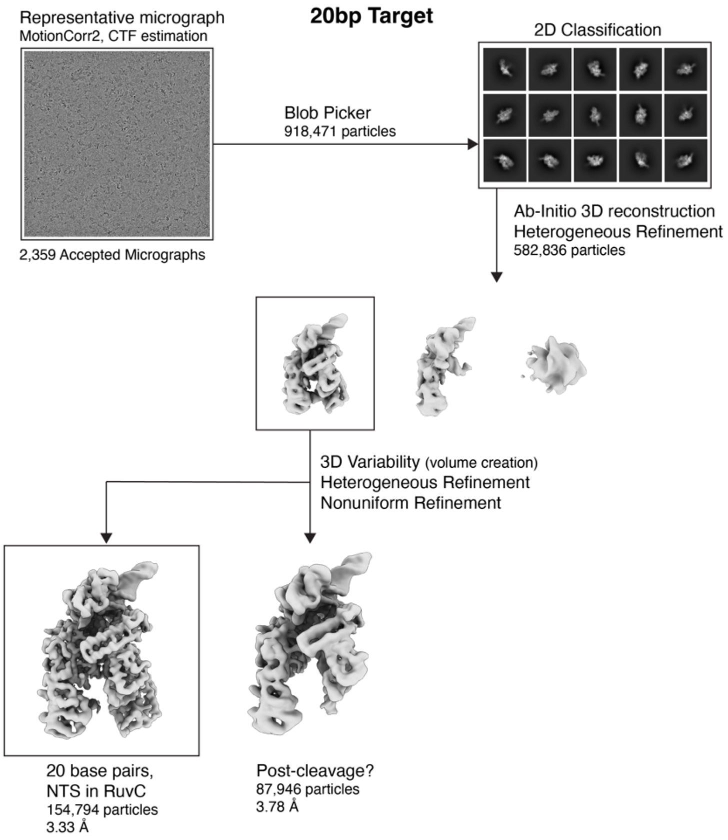


**Supp. Fig. 5 | Final data processing pipeline for 20bp target dataset.**

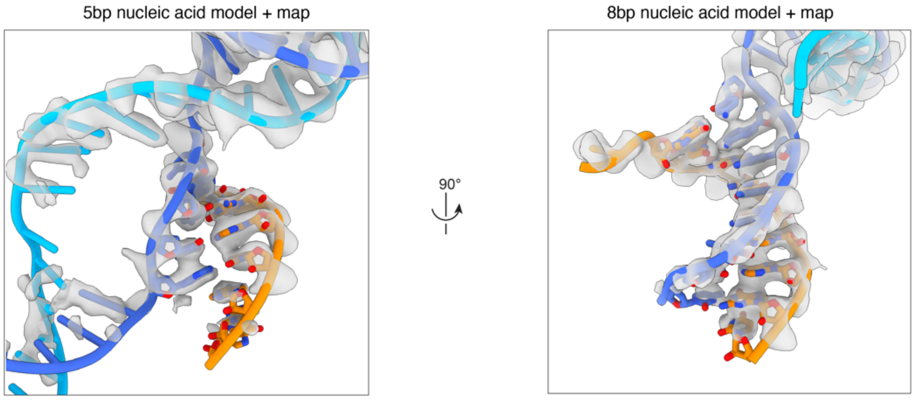


**Supp. Fig. 6 | Early R-loop structures within Cas12a.** Nucleic acid models from the early R-loop intermediates seen in Fig. 2 overlayed with their respective sharpened maps. The crRNA is shown excluding the pseudoknot and starting at position -1 (one base upstream of the guide sequence). It should be noted that in the 5bp reconstruction weak density can be observed for the target strand at position 6 of the R-loop, suggesting that a small fraction of particles base pair at this position.

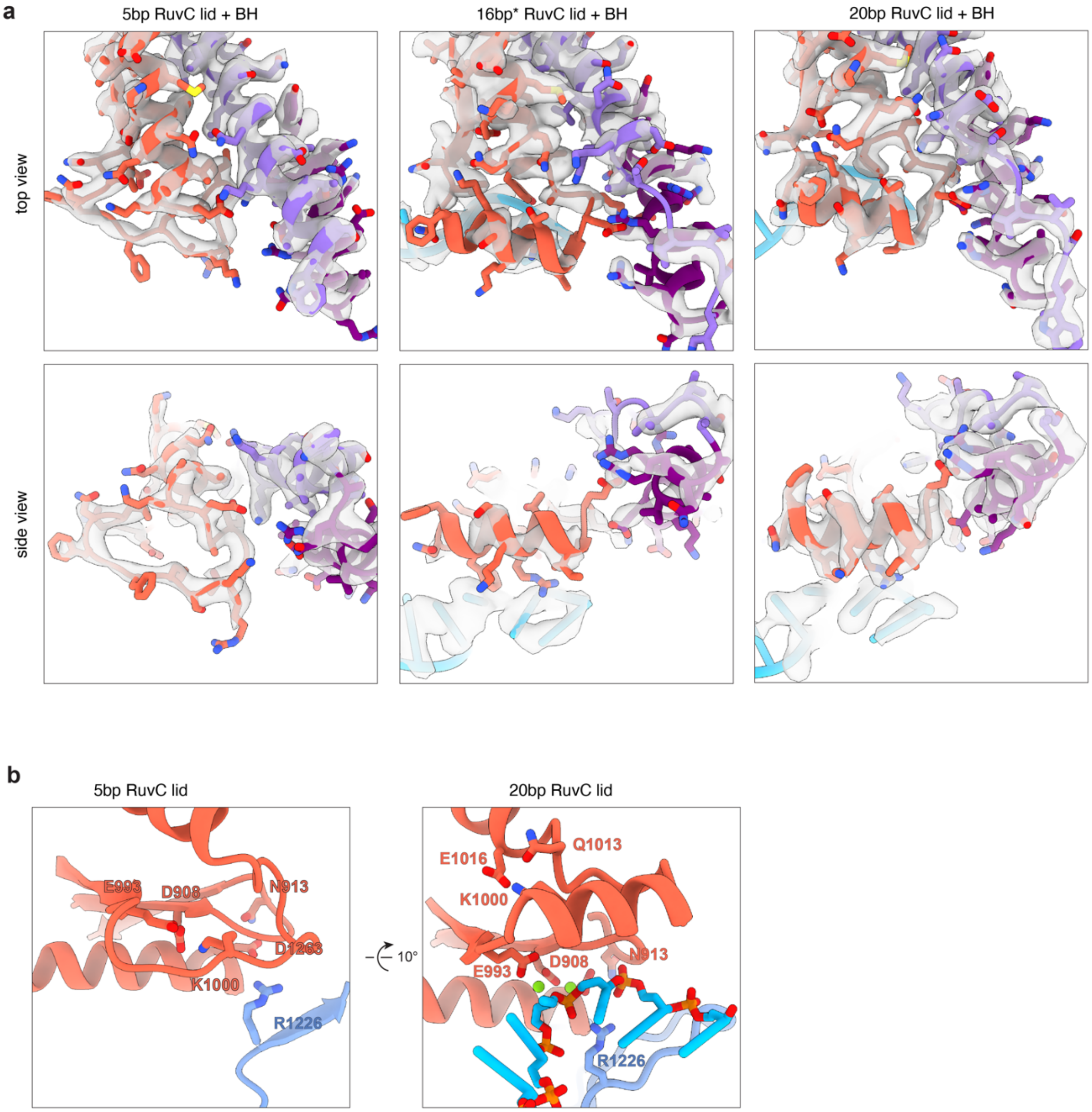


**Supp. Fig. 7 | Changes in RuvC lid conformation and contacts throughout R-loop formation. a,** Panel showing the model of the RuvC lid and BH domains overlayed with their respective sharpened maps, related to Fig. 3. Map density for the 16bp* structure is less well-resolved than the 20bp structure, suggesting structural heterogeneity due to lid movement or lack of helical structure. **b,** Comparison of the RuvC lid and active site contacts in the 5bp and 20bp structures.

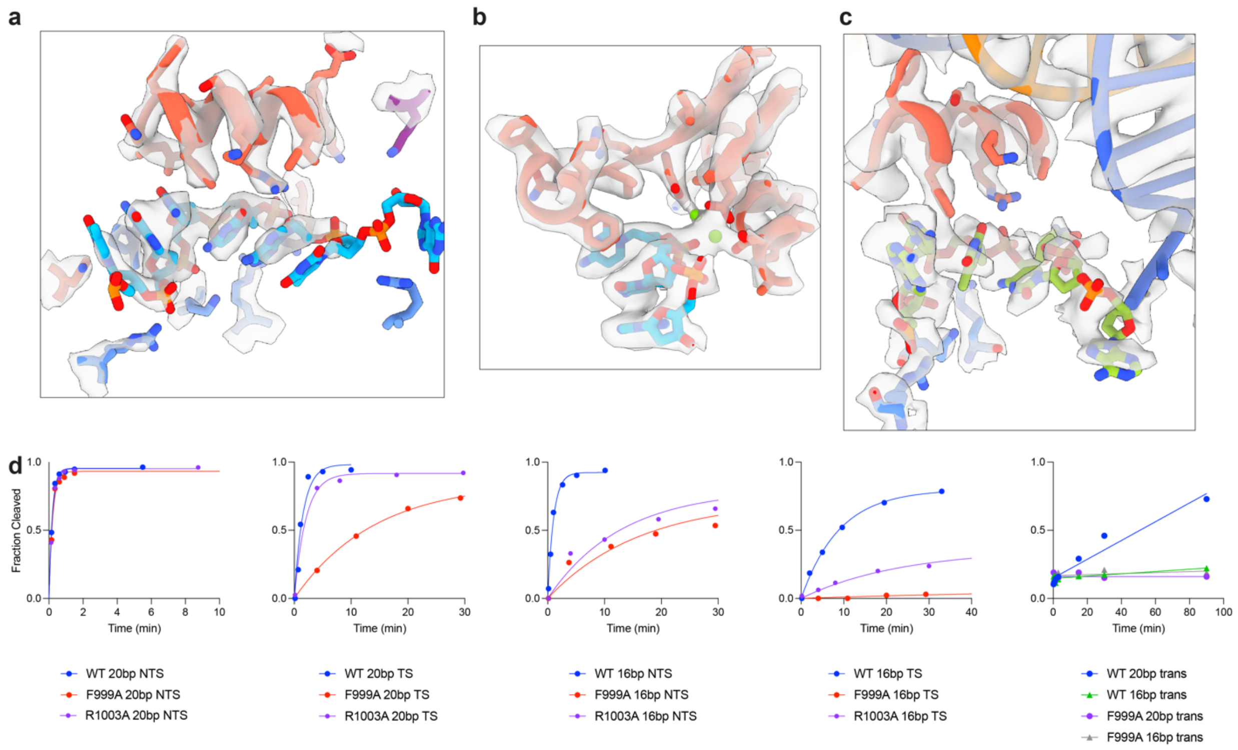


**Supp. Fig. 8 | Nucleic acids brought to the RuvC active site. a,** View of NTS in the RuvC active site, as in Fig. 4a, overlayed with sharpened map. **b,** View of NTS coordinated with the RuvC catalytic residues and $Mg^{2+}$ ions, as in Fig. 4b, overlayed with sharpened map. **c,** View of TS in the RuvC active site, as in Fig. 4f, overlayed with sharpened map. **d,** Example time courses of NTS, TS, and trans-cleavage by Cas12a and RuvC lid mutants F999A and R1003A.

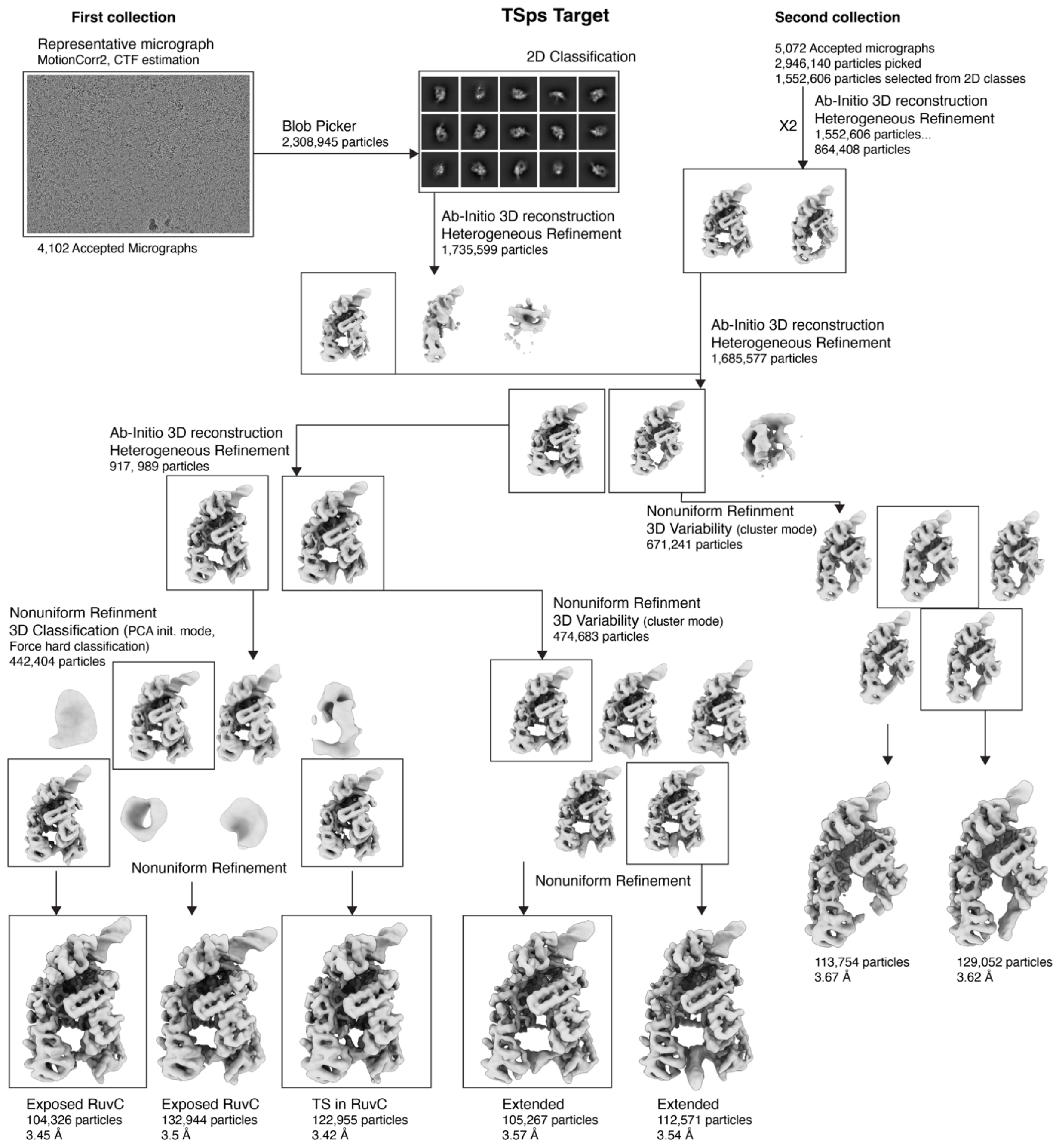


**Supp. Fig. 9 | Final data processing pipeline for TS (PS) target dataset.**

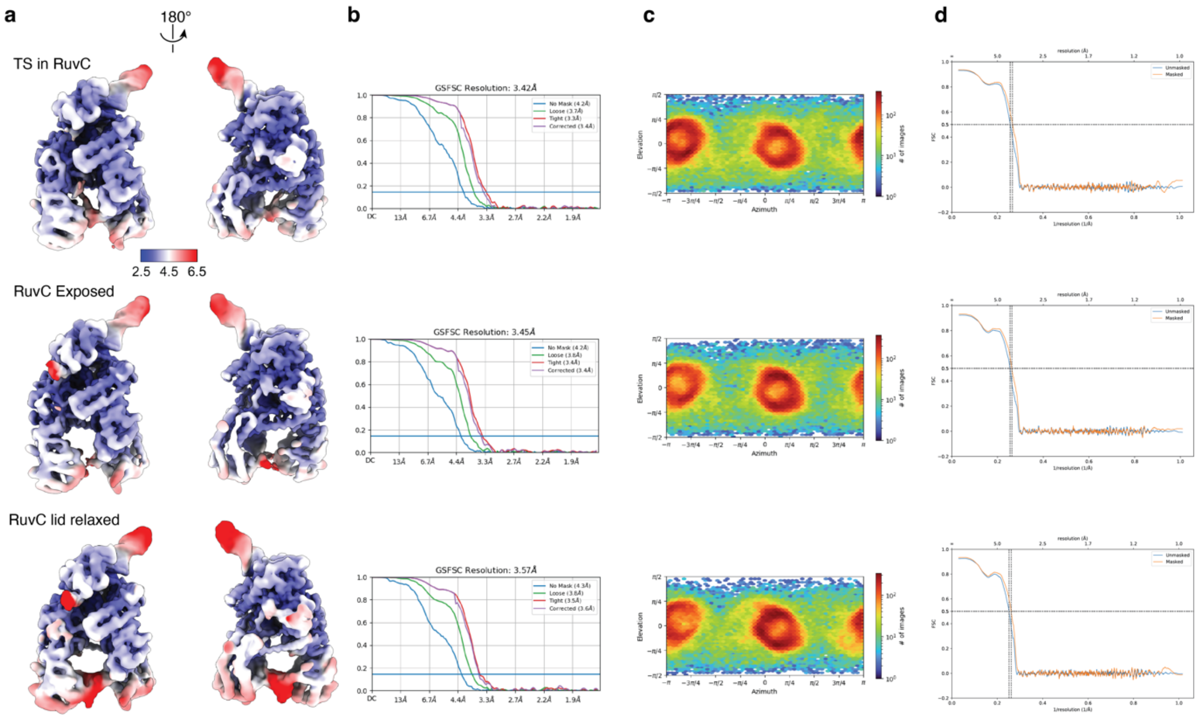


**Supp. Fig. 10 | Local resolution of TS(PS) maps.** **a,** Unsharpened maps of post-NTS cleavage reconstructions colored by local resolution. **b,** Gold-standard FSC curves for each reconstruction. Resolution estimated at FSC=0.143. **c,** Euler diagrams for each reconstruction showing particle orientation distributions. Map resolutions and particle distribution plots were produced in cryoSPARC. **d,** Map-to-model FSC curves produced in Phenix.

| | **5bp** (EMDB-40441) (PDB 8SFH) | **8bp** (EMDB-40442) (PDB 8SFI) | **10bp** (EMDB-40443) (PDB 8SFJ) | **15bp** (EMDB-40444) (PDB 8SFL) | **16bp*** (EMDB-40445) (PDB 8SFN) | **20bp** (EMDB-40446) (PDB 8SFO) | **TSps-TS in active site** (EMDB-40447) (PDB 8SFP) | **TSps-RuvC exposed** (EMDB-40448) (PDB 8SFQ) | **TSps-Extended** (EMDB-40449) (PDB 8SFR) |
|---|---|---|---|---|---|---|---|---|---|
| **Data collection and processing** | | | | | | | | | |
| Voltage (kV) | 300 | 300 | 300 | 300 | 300 | 200 | 300 | 300 | 300 |
| Electron exposure (e–/Å$^2$) | 80 | 80 | 80 | 80 | 80 | 49 | 80 | 80 | 80 |
| Defocus range ($\mu$m) | | | | | -1.5 to -2.5 | | | | |
| Pixel size (Å) | 0.833 | 0.833 | 0.833 | 0.833 | 0.833 | 0.94 | 0.833 | 0.833 | 0.833 |
| Symmetry imposed | | | | | C1 | | | | |
| Initial particle images (no.) | 1,624,766 | 1,624,766 | 1,509,051 | 3,758,114 | 3,758,114 | 582,836 | 3,288,205 | 3,288,205 | 3,288,205 |
| Final particle images (no.) | 60,595 | 47,461 | 122,723 | 158,948 | 123,714 | 154794 | 122,955 | 104,326 | 105,267 |
| Map resolution (Å) | 3.42 | 3.51 | 3.35 | 3.27 | 3.34 | 3.33 | 3.42 | 3.45 | 3.57 |
| FSC threshold | | | | | 0.143 | | | | |
| Map resolution range (Å) | | | | | 2.5 – 6.5 | | | | |
| **Refinement** | | | | | | | | | |
| Initial model used (PDB code) | | | | | | 5B43 | | | |
| Model resolution (Å) | 3.7 | 3.8 | 3.8 | 3.6 | 3.8 | 3.6 | 3.7 | 3.7 | 3.8 |
| FSC threshold | | | | | 0.5 | | | | |
| Map sharpening *B* factor (Å$^2$) | 132.2 | 128.5 | 173.6 | 162.8 | 147.5 | 154.4 | 171.7 | 165.7 | 172.4 |
| Model composition | | | | | | | | | |
| Non-hydrogen atoms | 12407 | 11477 | 10207 | 12973 | 12848 | 12606 | 12185 | 12045 | 12047 |
| Protein residues | 1286 | 1227 | 1001 | 1296 | 1302 | 1240 | 1240 | 1240 | 1240 |
| Nucleotides | 91 | 70 | 97 | 114 | 106 | 119 | 99 | 92 | 92 |
| Ligand | | | | | | MG: 2 | | | |
| Mean *B* factors (Å$^2$) | | | | | | | | | |
| Protein | 189.3 | 83.33 | 75.53 | 82.36 | 99.96 | 64.92 | 107.85 | 121.39 | 109.43 |
| Nucleotides | 265.32 | 106.3 | 129.35 | 182.19 | 137.78 | 105.18 | 125.13 | 122.06 | 115.71 |
| R.M.S. deviations | | | | | | | | | |
| Bond lengths (Å) | 0.004 | 0.005 | 0.005 | 0.005 | 0.005 | 0.004 | 0.004 | 0.005 | 0.005 |
| Bond angles (°) | 0.74 | 0.986 | 1.059 | 0.912 | 1 | 0.798 | 0.9 | 0.994 | 0.948 |
| Validation | | | | | | | | | |
| MolProbity score | 1.37 | 1.44 | 1.41 | 1.48 | 1.58 | 1.27 | 1.47 | 1.39 | 1.63 |
| Clashscore | 4.14 | 5.89 | 6.84 | 5 | 7.14 | 3.81 | 5.99 | 6.61 | 8.12 |
| Poor rotamers (%) | 0.61 | 0.55 | 0.34 | 0.69 | 0.6 | 0.18 | 0.63 | 0.72 | 0.63 |
| Ramachandran plot | | | | | | | | | |
| Favored (%) | 97.02 | 97.37 | 97.88 | 96.59 | 96.92 | 97.49 | 97.24 | 97.89 | 96.84 |
| Allowed (%) | 2.98 | 2.63 | 2.12 | 3.41 | 3.08 | 2.51 | 2.76 | 2.11 | 3.16 |
| Disallowed (%) | 0 | 0 | 0 | 0 | 0 | 0 | 0 | 0 | 0 |

**Supp. Table 1 | cryo-EM data collection and model stats.**

| Oligo | Sequence |
|---|---|
| D_min.crRNA | UUUUUAAUUUCUACUCUUGUAGAU**GUGAUAAGUGGAAUGCCAUG**UGGA |
| TargetD_NTS | CGCTCTTCCGATCTTTTA**GTGATAAGTGGAATGCCATG**TGGAGTAGCTACTGTGCT |
| TargetD_TS | AGCACAGTAGCTACTCCA**CATGGCATTCCACTTATCAC**TAAAAGATCGGAAGAGCG |
| 5comp_NTS | CGCTCTTCCGATCTTTTAGTGATTTCACCTTACGGTACTGGAGTAGCTACTGTGCT |
| 5comp_TS | AGCACAGTAGCTACTCCAGTACCGTAAGGTGAATTCACTAAAAGATCGGAAGAGCG |
| 8comp_NTS | CGCTCTTCCGATCTTTTAGTGATAAGACCTTACGGTACTGGAGTAGCTACTGTGCT |
| 8comp_TS | AGCACAGTAGCTACTCCAGTACCGTAAGGTCTTATCACTAAAAGATCGGAAGAGCG |
| 12comp_NTS | CGCTCTTCCGATCTTTTAGTGATAAGTGGATACGGTACTGGAGTAGCTACTGTGCT |
| 12comp_TS | AGCACAGTAGCTACTCCAGTACCGTATCCACTTATCACTAAAAGATCGGAAGAGCG |
| 16comp_NTS | CGCTCTTCCGATCTTTTAGTGATAAGTGGAATGCGTACTGGAGTAGCTACTGTGCT |
| 16comp_TS | AGCACAGTAGCTACTCCAGTACGCATTCCACTTATCACTAAAAGATCGGAAGAGCG |
| D_TSps | AGCACAGTAGCT*A*C*T*C*C*ACATGGCATTCCACTTATCACTAAAAGATCGGAAGAGCG |
| A4T_NTS | CGCTCTTCCGATCTTTTAGTGtTAAGTGGAATGCCATGTGGAGTAGCTACTGTGCT |
| A4T_TS | AGCACAGTAGCTACTCCACATGGCATTCCACTTAaCACTAAAAGATCGGAAGAGCG |
| ssD_20 | CATGGCATTCCACTTATCAC |
| ssD_16 | GCATTCCACTTATCAC |
| trans substrate1 | TTTATT |
| trans substrate2 | CCCCCCCCCC |

**Supp. Table 2 | List of oligos sequences used.** Sequences of RNA (Synthego) and DNA (IDT) used for sample preparation and biochemistry reactions. Oligos labeled for cleavage and binding detection had a 5'-FAM label added.

| Cas, R-loop | **NTS** ($min^{-1}$) | **TS** ($min^{-1}$) | **trans** ($M^{-1}$ $min^{-1}$) |
|---|---|---|---|
| WT, 20bp | 6.4 ± 1.2 | 0.77 ± 0.13 | 814000 ± 111000 |
| WT, 16bp | 1.5 ± 0.5 | 0.12 ± 0.01 | 38000 ± 5700 |
| F999A, 20bp | 4.5 ± 0.2 | 0.082 ± 0.011 | 350 |
| F999A, 16bp | 0.059 ± 0.003 | 0.0009 ± 0.0002 | 350 |
| R1003A, 20bp | 5.5 ± 0.8 | 0.55 ± 0.05 | - |
| R1003A, 16bp | 0.097 ± 0.019 | 0.018 ± 0.004 | - |
| LD, 20bp | 5.4 ± 0.4 | - | - |
| LD, A4T | 0.080 ± 0.016 | - | - |
| WT, A4T | 0.17 ± 0.02 | - | - |

**Supp. Table 3 | List of cleavage rates measured.** Cis-cleavage rates are listed as averages of at least duplicates ± standard error of the mean. Trans second order cleavage rates are listed as average of at least duplicates ± standard error of the mean.
